# Interdependence between confirmed and discarded cases of dengue, chikungunya and Zika viruses in Brazil: A multivariate time-series analysis

**DOI:** 10.1101/708743

**Authors:** Juliane F Oliveira, Moreno S. Rodrigues, Lacita M. Skalinski, Aline ES Santos, Larissa C. Costa, Luciana L. Cardim, Enny S. Paixão, Maria da Conceição N. Costa, Wanderson K. Oliveira, Maurício L. Barreto, Maria Glória Teixeira, Roberto F. S. Andrade

**Affiliations:** Center of Data and Knowledge Integration for Health (CIDACS), Instituto Gonçalo Moniz, Fundação Oswaldo Cruz, Salvador, Bahia, Brazil; Centre of Mathematics of the University of Porto (CMUP), Department of Mathematics, Porto, Portugal; Fundação Oswaldo Cruz, Porto Velho, Rondônia, Brazil; Universidade Estadual de Santa Cruz, Ilhéus, Bahia, Brazil; Instituto de Saúde Coletiva, Universidade Federal da Bahia, Salvador, Bahia, Brazil; London School of Hygiene and Tropical Medicine, London; Secretaria de Vigilância em Saúde, Ministério da Saúde, Brasília, Distrito Federal, Brazil; Instituto de Física, Universidade Federal da Bahia, Salvador, Bahia, Brazil

## Abstract

The co-circulation of different arboviruses in the same time and space poses a significant threat to public health given their rapid geographic dispersion and serious health, social, and economic impact. Therefore, it is crucial to have high quality of case registration to estimate the real impact of each arboviruses in the population. In this work, a Vector Autoregressive (VAR) model was developed to investigate the interrelationships between discarded and confirmed cases of dengue, chikungunya, and Zika in Brazil. We used data from the Brazilian National Notifiable Diseases Information System (SINAN) from 2010 to 2017. There were three peaks in the series of dengue notification in this period occurring in 2013, 2015 and in 2016. The series of reported cases of both Zika and chikungunya reached their peak in late 2015 and early 2016. The VAR model shows that the Zika series have a significant impact on the dengue series and vice versa, suggesting that several discarded and confirmed cases of dengue could actually have been cases of Zika. The model also suggests that the series of confirmed and discarded chikungunya cases are almost independent of the cases of Zika, however, affecting the series of dengue. In conclusion, co-circulation of arboviruses with similar symptoms could have lead to misdiagnosed diseases in the surveillance system. We argue that the routinely use of mathematical and statistical models in association with traditional symptom-surveillance could help to decrease such errors and to provide early indication of possible future outbreaks. These findings address the challenges regarding notification biases and shed new light on how to handle reported cases based only in clinical-epidemiological criteria when multiples arboviruses co-circulate in the same population.

**Author summary:** Arthropod-borne viruses (arboviruses) transmission is a growing health problem worldwide. The real epidemiological impact of the co-circulation of different arboviruses in the same urban spaces is a recent phenomenon and there are many issues to explore. One of them is the misclassification due to the scarce availability of confirmatory laboratory tests. This establishes a challenge to identify, distinguish and estimate the number of infections when different arboviruses co-circulate. We propose the use of multivariate time series analysis to understand how the weekly notification of suspected cases of dengue, chikungunya and Zika, in Brazil, affected each other. Our results suggest that the series of Zika significantly impact on the series of dengue and vice versa, indicating that several discarded and confirmed cases of dengue might actually have been Zika cases. The results also suggest that the series of confirmed cases of chikungunya are almost independent of those of dengue and Zika. Our findings shed light on yet hidden aspects on the co-circulation of these three viruses based on reported cases. We believe the present work provides a new perspective on the longitudinal analysis of arboviruses transmission and call attention to the challenge in dealing with biases in case notifications when multiple arboviruses circulate in the same urban environment.

## Introduction

In recent times, the re-emergence and the rapid spread of arboviruses in urban areas have become a serious problem that has concerned health authorities as well as the general population in many countries. The magnitude of the epidemics, the occurrence of severe cases with neurological manifestations and lethal outcomes, and severity of congenital malformations associated with infections occurred during pregnancy are the main threats of this new epidemiological situation [1, 2].

In Brazil, the co-circulation of the four serotypes of dengue virus (DENV), together with the emergence and dissemination of chikungunya virus (CHIKV) and Zika virus (ZIKV), transmitted by *Aedes* mosquitoes (mainly *Aedes aegypti*), has a relevant negative impact on the health of the population and lead to an increase in the demand on health and other support services. From the DENV introduction, in 1986, to until arrival and subsequent spread of CHIKV and ZIKV, dengue was the most important vector-borne disease circulating in cities of Brazil, [3, 4]. In September of 2014, CHIKV was detected in cities of the states of Amapá and Bahia. Although chikungunya causes arthralgia with pain at a higher level than dengue, the other symptoms are similar, which increased the likelihood of misdiagnosis [5]. In October 2014, an outbreak of an undetermined exanthematous illness was registered in Rio Grande do Norte, in the northeast of Brazil, while in April 2015, ZIKV was identified as its aetiologic agent [6]. Patients infected with ZIKV typically presented low (or no) fever and skin rash within the first 24 hours of the disease onset, while DENV and CHIKV cause high fever immediately. Also, CHIKV causes more intense arthralgia than DENV. However, the other symptoms are similar, which confound and complicate their differentiation and easily lead to misdiagnosis, [2, 7, 8].

The similarity of symptomatology has made the clinical diagnosis of arboviruses difficult, especially in the course of epidemics with viral co-circulation, in which laboratory tests are still unavailable for most patients. The misclassification and incorrect diagnosis affect the risk estimates of these diseases, since epidemiological surveillance depends on the quality of the data to provide morbidity and mortality information close to the reality lived by the population and, consequently, the development of effective prevention strategies, [2, 7, 9]. Therefore, this study aims to explore and understand how dengue notified cases were impacted by the introduction and spread of chikungunya and Zika virus in Brazil.

## Materials and methods

We used a multivariate time series analysis in order to understand how the classification of notified cases of dengue, chikungunya and Zika affected each other in Brazil from 2015 to 2017.

### Data source

We used data from the Brazilian National Notifiable Diseases Information System (SINAN). We collected weekly reported data of suspected cases of dengue (from 2010 to 2017), chikungunya (from 2014 up to 2017), and Zika (from 2015 up to 2017) viruses. We only considered cases that presented non-null information about the temporal variable, i.e., week of notification or week of first symptoms. We further separated the cases into confirmed and discarded, following the final classification information as used by the Brazilian Ministry of Health. Confirmed cases are all suspected cases of disease, excluding those discarded or inconclusive. This classification can be based on clinical/epidemiological criteria, namely presence of clinical symptoms in the same area and time as other probable cases, or on clinical/laboratory criteria, namely the presence of clinical symptoms and a positive IgM ELISA result, viral RNA detection via PCR, NS1 viral antigen detection, or positive viral culture [3]. Discarded cases are defined as any suspected case that satisfies at least one of the following criteria: negative laboratory diagnosis (IgM serology); a laboratory confirmation of another disease; clinical and epidemiological compatibility with other diseases. Inconclusive cases of dengue and chikungunya (≲ 10% reported cases) were excluded from the analyses. However inconclusive cases of Zika represented about 30% (110,656/361,396 registered cases) of all notified cases from 2015 to 2017, accounting for about 56% (33,863/60,972 registered cases) of the Zika notifications in 2015. Once specific Zika laboratory tests were unavailable in that period, for the purpose of the carried analyses, we considered inconclusive Zika cases as confirmed Zika cases.

To perform the study of time series analyses, we collected the confirmed and discarded reported cases of each arbovirus per epidemiological week in Brazil, from 2015 to 2017.

### Multivariate time-series analysis

We constructed a Vector Autoregressive (VAR) model to uncover possible correlation and causality effects between the discarded and confirmed cases of the three arboviruses.

Formally, a time-series is defined as *random sequence*, i.e., a collection of random variables {*Y_t_*}, where the time-index assumes integer values only. A univariate time-series {*Y_t_*} is said to be an *autoregressive process* if each *Y_t_* is defined in terms of its predecessors *Y_p_*, for *p* < *t*, by the equation:

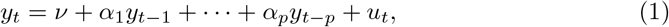

where *ν* is a fixed intercept constant allowing to the possibility of a non-zero mean, the *α_i_* are the linear coefficients, and {*U_t_*} is a *white noise*, i.e., a sequence of mutually independent random variables, each with mean zero and finite variance *σ*^2^.

If a *m*-dimensional multivariate time-series {***Y**_t_* = (*Y*_1*t*_, ⋯, *Y_mt_*)} is considered, a VAR process is defined as a generalization of equation (1), where *ν* is replaced by a constant *m*-vector ***ν*** of components *ν_i_*, the linear coefficients *α_i_* are replaced by (*m* × *m*)-matrices ***α^i^*** of elements 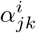, and ***U_t_*** is a multivariate *m*-vector of white noise components *u_it_*. Therefore, the generalized form of equation (1) in matrix form is:

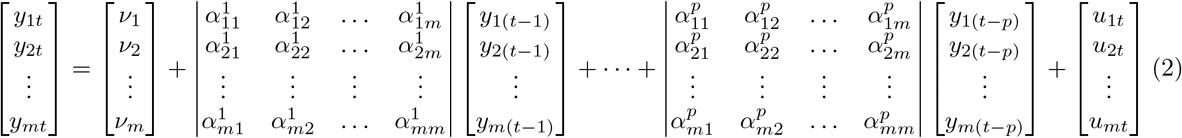

We say that equation (2) is a VAR process of order *p*, VAR(*p*), if ***α***^1^, ⋯, ***α**^p^* ≠ 0 and ***α**^i^* = 0 for *i* > *p*, where *p* is the smallest possible order.

In this work, we considered a 6-dimensional multivariate time series, with each *Y_t_* representing a vector (*Z*_1*t*_, *Z*_2*t*_, *C*_1*t*_, *C*_2*t*_, *D*_1*t*_, *D*_2*t*_), where *Z*, *D*, and *C* denotes Zika, chikungunya and dengue respectively, the indices 1 and 2 stand for confirmed and discarded cases, and the time *t* ranges from the first week of 2015 until the last week of 2017.

In order to carry out the analysis, we first transformed the vector series {*Y_t_*} to a stationary form, in such a way that its mean value and the covariance among its observations do not change with time. The used transformation is given by:

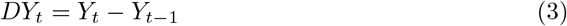

After performing the transformation, the Dickey-Fuller Test was used to certify that the resulted series are indeed stationary.

Using equation (2), the stationary series (3) were written in terms of its own terms and of the other series on *p* previous weeks. The order (lag) *p* was selected by the Akaike information criterion (AIC), which is based on the minimization of the the mean squared error of complete set of adjusted series [10].

The steps to construct and analyse a VAR(p) model amounts to: (i) estimate the VAR coefficients by a multivariate Least Squares Estimation; (ii) select the *p* (lag) order using AIC; (iii) test for normality of residuals by representing their ordered values as a function of theoretical quantiles, by probability plots that show how the they depart from normal curve, and by the analysis of the partial (cross-)correlation function (PCF) between them; (iv) perform pair-wise Granger tests among the 6-dimensional estimated multivariate series.

Using the Granger causality F-test in a pair-wise way, we can check whether the null hypothesis, stating that one series {*X_t_*} does not affect the other one {*Y_t_*}, can be rejected or not. If the hypothesis is rejected, then the time-series {*X_t_*} affects the series {*Y_t_*}. Thus, from a rather heuristic point of view, this indicates that past values of {*X_t_*} can be used for the prediction of future values of {*Y_t_*}. Otherwise stated, the values used for describing the autoregressive equation for {*Y_t_*} have significant non-zero contribution of past values of {*X_t_*}. The model including {*X_t_*} is called unrestricted model, in opposition to the restricted case where the series {*X_t_*} is not included in the analysis, see [11].

Detailed information about the theoretical background for time series analysis can be found in [10, 12]. We performed our statistical analysis using a specifically developed Python code based on the Statsmodels Tools [13, 14].

### Ethics Statements

This is an ecological study conducted with anonym secondary data of public domain and therefore does not require approval of Human Research Ethics Committee/HREC. However, we submitted it to the HREC of the Federal University of Bahia (Salvador/BA) and we obtained ethics approval - CAAE: 70745617.2.0000.5030.

## Results

From January 2010 to December 2017 were registered in SINAN 12,130,550 million cases of dengue, from which 52% were confirmed and 32% were discarded. From January 2014 to December 2017 were registered in SINAN 501,202 thousands of cases of chikungunya, resulting in 63% and 27% of confirmed and discarded cases, respectively. The classified confirmed and discarded cases of Zika, from January 2015 to 2017, represented 76% and 24% of registered cases, respectively.

During the period covered by this study, the confirmed cases of dengue had three main peaks that occurred in 2013 (with 1,189,370 registered confirmed cases), in 2015 (with 1,389,784 registered confirmed cases) and in 2016 (with 1,101,228 registered confirmed cases). Confirmed Zika and chikungunya cases reached their peaks in late 2015 and middle of 2016, respectively. Fig 1 shows, in a single panel, the evolution of curves of confirmed and discarded cases in the time window where the three diseases occurred simultaneously (2015 - 2017).

**Fig 1.**
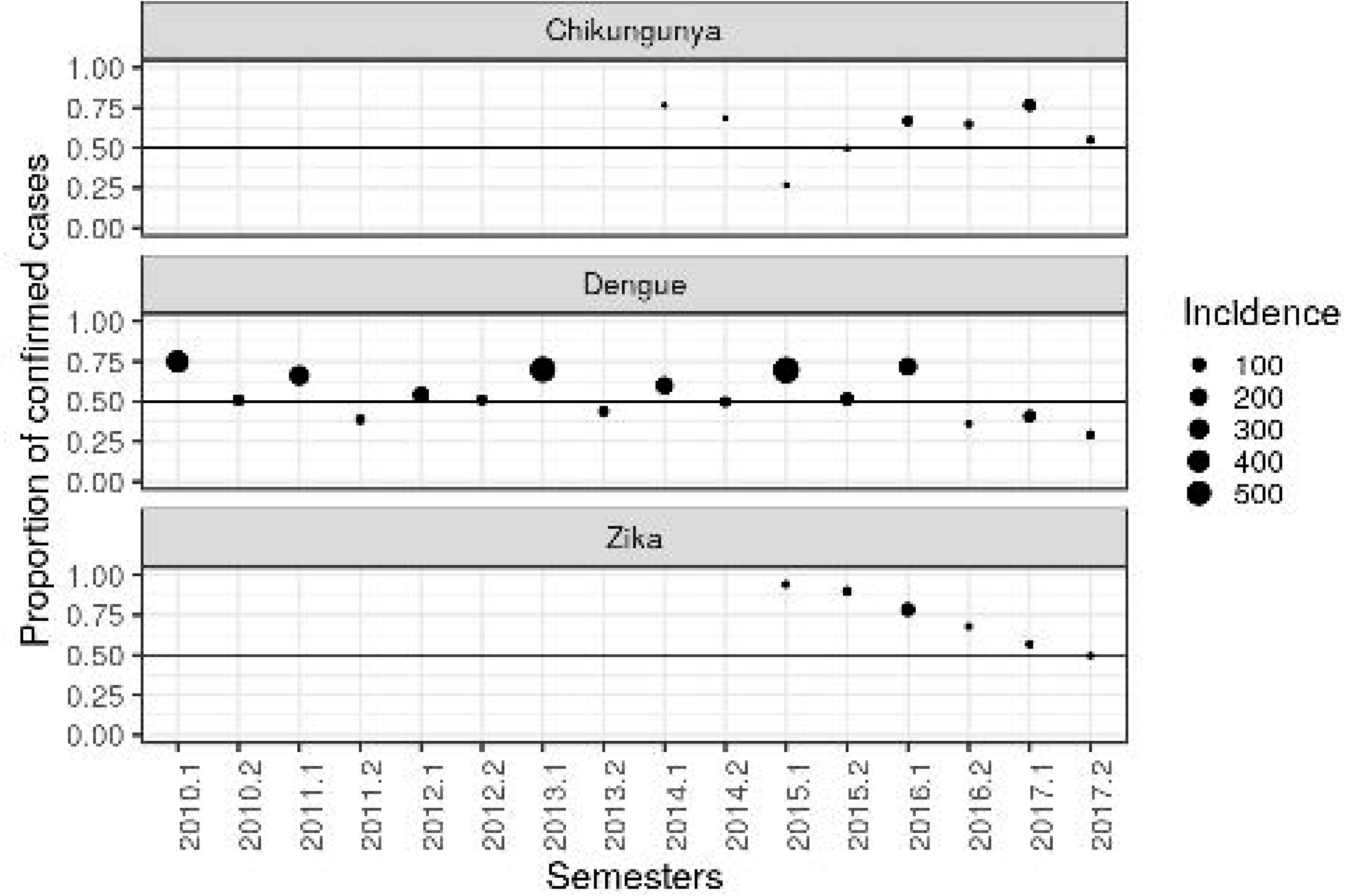
Illustration of time series plots of confirmed and discarded cases of dengue, chikungunya and Zika by epidemiological week. Brazil, January 2015 to December 2017.

Exploring the linear dependence between the series of confirmed and discarded cases, per epidemiological week, during the whole corresponding periods, for dengue, chikungunya and Zika in the country, we see that the slope *a_D_* of the linear relation for dengue is 2.0, while yearly based evaluations lead to a mean value 〈*a_D_*〉_*t*_ = 2.3 (*SD* = 1). It means that, on average, for every ten confirmed cases of dengue, about 4 cases are discarded. For chikungunya and Zika, the slopes are 3.2 and 3.6 respectively.

In Fig 2, we plot the ratio between the number of confirmed cases and the sum of confirmed and discarded cases, per semester. Given the well documented seasonal behaviour of the dengue epidemics during the last three decades, we adopt the used identification of the first and second semester of each year as the epidemic and inter-epidemic periods, respectively. The closer to 0.5 the dots are, the number of confirmed and discarded cases are close to each other.

**Fig 2.**
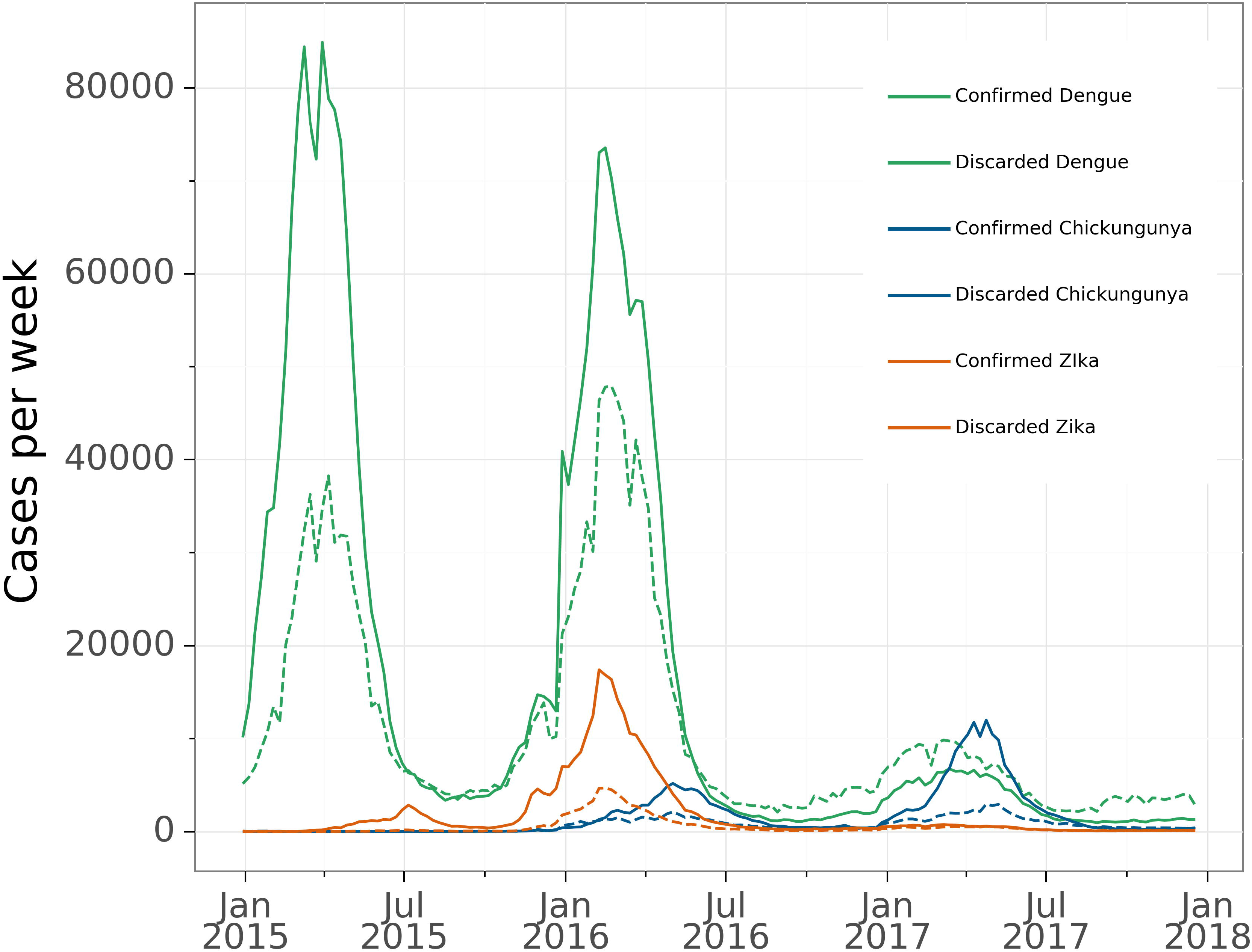
**Proportion of confirmed and discarded cases, per semester, of dengue, chikungunya and Zika in Brazil** proportions of dengue (from 2010 up to 2017); chikungunya (from 2014 up to 2017); Zika (from 2015 up to 2017).

We can see that the patterns of the proportion between confirmed and discarded cases differ among the diseases. For dengue, the proportion of discarded cases are most common during the second semester of the year, that is, in the inter-epidemic period. This pattern is also observed in years with the lowest incidence of dengue, namely 2012, 2014 and 2017.

Although chikungunya presents lower incidence compared with Zika and dengue, in 2015 we notice an inverted pattern in the proportion of confirmed and discarded cases compared to those reported by dengue and Zika. In 2017, where the proportion of discarded cases is higher for dengue and Zika, chikungunya shows a pattern with a higher proportion of confirmed cases in the first semester and lower in the second. The proportions of confirmed cases are more common both in the first and second semesters.

The first step to proceed with the multivariate time-series analysis is by checking that the series to be considered are stationary. Using the Dickey-Fuller test applied for each of the series of confirmed and discarded cases of dengue, chikungunya and Zika, the results shows the series are stationary after one differentiation (all presenting p-values < 0.005). S1 Table shows a summary of the results of the applied test to the considered six series before and after differentiation.

The empirical determination of the appropriate lag order considered all six series (Z,D,C discarded and confirmed), for which the lag *p* according to the AIC is 13, see S2 Table. Table 1 shows the correlation matrix of the stationary series of confirmed and discarded cases of dengue, chikungunya and Zika. We interpret the values of positive/negative correlation according to the interval: ±.00 to ±.10 very low; ±.10 to ±.40 as weak; ±.40 to ±.60 as moderate; ±.60 to ±.80 as strong; ±.80 to ±.99 very strong; ±1.0 as perfect. Summary of regression results are presented in S1 Appendix. The autocorrelation, cross-correlations and probability plots of residuals are given from S1 Fig to S4 Fig.

**Table 1.** Correlation matrix of the stationary series of confirmed and discarded cases of dengue, chikungunya and Zika. Brazil, January 2015 to December 2017.

|  | *Conf.Zika* | *Disc.Zika* | *Conf.chik.* | *Disc.chik.* | *Conf.dengue* | *Disc.dengue* |
| --- | --- | --- | --- | --- | --- | --- |
| *Conf.Zika* |  | .93 | .04 | .17 | .63 | .70 |
| *Disc.Zika* | .93 |  | .10 | .27 | .66 | .75 |
| *Conf.chik.* | .04 | .10 |  | .66 | .03 | .04 |
| *Disc.chik.* | .17 | .27 | .66 |  | .23 | .23 |
| *Conf.dengue* | .63 | .66 | .03 | .23 |  | .69 |
| *Disc.dengue* | .70 | .75 | .04 | .23 | .69 |  |

p-values < 0.005). S1 Table shows a summary of the results of the applied test to the considered six series before and after differentiation.

The results of the Granger tests to explore associations between series are presented in Table 2. They show that, at a significant level, the series of confirmed and discarded cases of dengue affect the series of confirmed and discarded cases of Zika and vice-versa. On the other hand, no associations were found between the series of confirmed and discaded cases of Zika and chikungunya. The series of confirmed Zika and confirmed (0.63) and discarded (0.70) dengue, as well as, the series of discarded Zika and confirmed (0.66) dengue present a positive strong linear correlation, which is stronger than the other correlations for the series described above.

**Table 2.** Results of pairwise Granger tests. Exploratory search of associations between series of confirmed and discarded cases of dengue, chikungunya and Zika. Brazil, January 2015 to December 2017.

| Null hypothesis: | Test statistic | p-value | Result |
| --- | --- | --- | --- |
| Confirmed cases of dengue do not affect confirmed cases of Zika | 3.836 | < 0.001 | Reject |
| Confirmed cases of Zika do not affect confirmed cases of dengue | 5.363 | < 0.001 | Reject |
| Discarded cases of dengue do not affect confirmed cases of Zika | 3.836 | < 0.001 | Reject |
| Confirmed cases of Zika do not affect discarded cases of dengue | 4.112 | < 0.001 | Reject |
| Confirmed cases of dengue do not affect discarded cases of Zika | 4.567 | < 0.001 | Reject |
| Discarded cases of Zika do not affect confirmed cases of dengue | 3.417 | < 0.001 | Reject |
| Confirmed cases of chikungunya do not affect confirmed cases of dengue | 2.121 | 0.012 | Reject |
| Confirmed cases of chikungunya do not affect discarded cases of dengue | 1.942 | 0.025 | Reject |
| Discarded cases of chikungunya do not affect confirmed cases of dengue | 2.172 | 0.010 | Reject |
| Confirmed cases of dengue do not affect confirmed cases of chikungunya | 0.4283 | 0.959 | Do not reject |
| Confirmed cases of dengue do not affect discarded cases of chikungunya | 0.6629 | 0.799 | Do not reject |
| Discarded cases of dengue do not affect confirmed cases of chikungunya | 1.070 | 0.384 | Do not reject |
| Confirmed cases of chikungunya do not affect confirmed cases of Zika | 1.128 | 0.333 | Do not reject |
| Confirmed cases of Zika do not affect confirmed chikungunya | 0.9064 | 0.547 | Do not reject |
| Discarded cases of chikungunya do not affect confirmed cases of Zika | 0.9579 | 0.493 | Do not reject |
| Confirmed cases of Zika do not affect discarded cases of chikungunya | 0.9339 | 0.518 | Do not reject |
| Confirmed cases of chikungunya do not affect discarded cases of Zika | 1.010 | 0.440 | Do not reject |
| Discarded cases of Zika do not affect confirmed cases of chikungunya | 0.8859 | 0.568 | Do not reject |

There is a significant association between confirmed cases of chinkugunya and confirmed and discarded cases of dengue. However, by assessing the correlation matrix given in Table 1, the values present a very low correlation (0.03 and 0.04, respectively). We also found that discarded cases of chikungunya have a significant association with the series of confirmed cases of dengue. This last showing a positive weak correlation (0.23).

In S2 Table we present AIC values for the unrestricted model involving the six series and the restricted ones, where by turn confirmed and discarded cases of either dengue or chikungunya or Zika are not included. Additionally, we show the results of the respective Granger causality tests in S3 Table.

The restricted model excluding the series of chikungunya has a better performance (AIC = 48.50, lag order = 13) regarding the multi-series model including the six series. An analysis of the residuals and model fitting similar to what is presented here was also done, again showing better results. The Granger tests performed for these four series also stay in agreement with the results for the unrestricted model including the six series.

Although it is clear a mutual influence of the cases of Zika and dengue, we also performed an analysis of the restricted models between the cases of Zika and chikungunya and the cases of dengue and chikungunya. The analysis between Zika and chikungunya also presents a better AIC value 41.67 (lag order = 13). However, the Granger tests shows three different results: i) confirmed cases of Zika affect discarded cases of chikungunya (Test statistic = 2.206, p-value = 0.011); ii) discarded cases of Zika affect confirmed cases of chikungunya (Test statistic = 2.447, p-value = 0.004); and iii) a borderline result where confirmed cases of Zika affect confirmed cases of chikungunya (Test statistic = 1.761, p-value = 0.053). To conclude, the Granger tests for the analysis between dengue and chikungunya (AIC = 52.06, lag order = 12) presented less relations among them, with only confirmed cases of chikungunya affecting discarded cases of dengue. Despite this, in both models, a visual analysis through the probability plot of the residuals does not suggest normality.

## Discussion

In the context of mutual influence of notifications of co-occurring diseases, a series of discarded cases of disease 1 may affect the confirmed cases of disease 2. In such cases, it is possible to argue that individuals truly infected by disease 2 were wrongly notified as disease 1 provided there exists strong similarities and weak differences among main signs and symptoms of each disease, as it has been reported since 1969 for arboviroses [3, 15, 16]. On the other hand, when confirmed cases of disease 1 affect the discarded cases of disease 2, this can be interpreted as an increase (or decrease) in the notification of disease 2, but not necessarily this notification can be claimed as a confirmed case of disease 1.

The strong positive correlation found in the analyses shows that the notification series of dengue was significantly impacted by Zika, and vice-versa. A reasonable interpretation is that an increase of individuals notified as Zika contributes to an increase of wrong notification of dengue cases within two scenario: people with Zika were wrongly classified as dengue (and vice-versa), or perhaps both viruses infected the same individuals. The results also indicate that the series of confirmed Zika cases increased the series of discarded dengue cases and that, among those discarded, there were indeed confirmed Zika cases. The series of confirmed cases of dengue affects significantly and positively (increasing) the series of discarded cases of Zika, which can be explained by the awareness of the consequences accounted by Zika at the end of 2015.

The notifications of Zika and dengue had overall weaker effects on the notification of suspected chikungunya cases, as indicated by the smaller correlation values between the corresponding series. The association of these values with Granger scores indicates a possible influence in 4 out of the 12 possible combinations. Considering confirmed and discarded cases of Zika and chikungunya, the unrestricted model was not able to find associations between these series. The analyses suggest that they did not affect each other and probably their notifications happened as independent events in Brazil. As to the mutual influence of dengue and chikungunya notifications, Granger scores show that possibly a small amount of confirmed and discarded cases of dengue were either confirmed cases of chikungunya or dengue notification were increased by the notification of chikungunya. In opposition to these specific combinations, the remaining results show either the failure to reject the null hypothesis, as in the result for chikungunya and Zika cases, or a very small correlation value, as for both confirmed dengue and chikungunya cases. Thus, the remaining they also support the conclusion that the notifications of dengue and Zika happened independently of the notification of chikungunya.

Regarding the proportions of reported confirmed and discarded cases of each viruses, although chikungunya presents lower incidence compared with Zika and dengue, in 2015 we notice an inverted pattern in the proportion of confirmed and discarded cases compared to those reported by dengue and Zika. This can rise up the hypothesis that one virus inhibits the proliferation of the other. In 2017, when the proportion of discarded cases is larger for dengue and Zika, chikungunya shows a pattern with a higher proportion of confirmed cases in the first semester and lower in the second. Ribeiro et al [17], performed surveillance study in the city of Salvador, Brazil. They argued that the increased pattern of chikungunya cases in opposition to the decrease of dengue and Zika cases in this population, may suggest that it was unrelated to vector population and that immunity after ZIKAV infection may cross-protect against dengue. Our findings, using data of the whole Brazil, are in accordance with their results.

This study highlights that in Brazil, from 2015 to 2017, the series of notifications of confirmed and discarded cases of dengue, chikungunya and Zika presented, in most of the cases, linear dependence. This reflects the epidemiological context presented by this country from the second semester of 2014 on, when the simultaneous circulation of DENV, ZIKV and CHIKV in densely populated urban spaces greatly hampered the correct record of each case of these diseases [9, 18]. Although CHIKV and ZIKV emerged almost simultaneously in cities in the same region of the country, the latter was only identified at the end of April 2015 [6]. Thus, there was a delay in alerting the health services network about the existence of this new clinical entity. In spite of the long experience in dengue of the professionals of the network of health services of this country, the circumstances presented above did not allow the adequate clinical and epidemiological diagnosis of the cases of each of these three diseases, resulting, often than not, in incorrect records [2].

In a scenario where only the notification of the diseases are available and laboratory tests are scarce, we see that the notification of dengue and Zika are shown to be independent of the notification of chikungunya. These findings are plausible, since dengue and Zika present more similar clinical manifestations to each other as compared to chikungunya [7]. The expressive joint manifestations produced by CHIKV infections allow a more accurate clinical diagnosis, even when specific laboratory tests are not available. Therefore, these results would not support the use of discarded cases of chikungunya as complementary cases of Zika infection, as suggested by Oliveira et al [18]. However, as the total number of chikungunya discarded cases was small (3.8 %) in comparison to the universe of cases of the three diseases, that fact did not affect the temporal trend presented for this and our study. Nevertheless, it is worth noticing that our results should be nearest of the real, and thus contribute to construct more accurate prediction models of future Zika epidemics, using only possible cases of dengue.

Another point that calls attention is that the lack of association between the series of confirmed cases of dengue and confirmed chikungunya, and confirmed cases of Zika and confirmed chikungunya might be due to spatial factors that are not considered in this work, or to a hypothetical situation where one virus inhibits the proliferation of the other.

Our studies based on a rather simplified linear model can be complemented by future works, where the analysis proceeds either through non-linear methods or through a more comprehensive and adequate model. A detailed study, probably based on the symptoms presented by patients, may also contribute to having better estimations for the quantity of cases that can be assigned for each disease. All suppositions made here are based only on a temporal analysis of the time series of notifications. Therefore, including a spatial analysis would clarify more issues regarding the surveillance of co-circulation of arboviruses and how zones with higher incidences handled with the notifications of the cases.

According to Barbosa et al (2015), at least until 2010, the dengue surveillance system in Brazil was able to inform about the temporal and spatial trends of the occurrence of this disease, providing good evaluation indicators [19]. However, with the emergence of CHIKV and ZIKAV in the period 2014-2015, which have clinical manifestations similar to those of dengue, and in a scenario of limited availability of laboratory tests, there were major difficulties with this system. From this time on, the diagnosis of the viruses were mostly the clinical and epidemiological, limiting the correct registration of the cases in the SINAN. As it is of crucial importance for the surveillance to know, as close as possible, the real magnitude of occurrence of each of these arboviruses, it is considered that this study provided important issues that can contribute to improve the risk estimate and the spatial dissemination knowledge of the diseases. At this stage of understanding, we believe that our results raise a discussion of misregistration and suggest directions to overcome such a difficulty.

These three relevant public health problems produce a high burden of disease for the population, as well as giving more complexity to the disease notification system, case by case, due to the similarity of the clinical features in the acute phase of these arboviruses, which are transmitted by the same vectors and in the same population spaces. Thus, there is no doubt that the actions of health services for its prevention and control should be developed in an integrated manner. However, it is necessary that the public health surveillance system seeks to improve its specific diagnostic strategies, not only to know the epidemiological profile in each space, but especially to enable appropriate clinical management in the acute form, as the cases with suspected dengue that need more massive and immediate hydration to prevent deaths; when it comes to Zika, patients should be warned of the possibility of post-acute neurological forms requiring hospitalization, as well as special attention to pregnant women and guidance for women of childbearing age to protect themselves from insect bites and sexual transmission; as for chikungunya, an important proportion of cases require monitoring and management of joint pain. Thus, improving case definitions requires conducting validation studies as well as training health professionals involved in patient care in order to make them better able to make diagnostic suspicions. It is essential to expand the offer of specific laboratory tests, but nowadays such tests for acute phase of the disease are almost restricted to molecular tests (due to the similarity of antibodies against dengue and Zika), which are very expensive to use on a large scale. In this sense, studies focused on the development, with different technologies, of sensitive and specific serological tests for these two flaviviruses, are a good prospect to be used in the routine of health services, as they are cheaper and easier to use. It is understood that, alongside the integration of surveillance of urban arboviruses, these initiatives should be adopted to overcome the difficulties of clinical management, such as the production of more reliable epidemiological data, which are of the greatest relevance for the planning of public health actions and for the prevention of these diseases.

In summary, we demonstrate two important interrelated aspects: the first one refers to how the discarded cases, which resulted from reported cases of one arbovirus, can be considered as part of complementary notifications of another; the second concerns how the series of confirmed cases of one disease may affect the series of confirmed cases of another. Thus, these findings address the challenges regarding notification biases and shed new light on how to handle reported cases based only in criteria clinical-epidemiologic when these three arboviruses co-circulate in the same population. We would like to emphasize that we could not find in the literature similar results aiming to discuss or explore large sets of occurrences where inter-correlations of cases discarded from one arbovirus and confirmed in the other could exist. A possible exception is a small scale, clinically based study concerned to the definition of cases, whereby the analyses were performed by checking if the symptoms presented by patients, after a second analysis by trained specialists, were well associated to one specific arbovirus [20]. In view of this, the results presented in this article may constitute one of the first attempts to systematically address this issue.

## Supporting information

Results of Dickey-Fuller Test applied to the series of confirmed and discarded cases of dengue, chikungunya and Zika before and after differentiation.

AIC and Lag values for the restricts and unrestricted models for the series confirmed and discarded cases of dengue, chikungunya and Zika. Brazil, Jan

Results of pairwise Granger tests for restricted models.

Summary of regression results

## Supporting information

**S1 Table. Results of Dickey-Fuller Test applied to the series of confirmed and discarded cases of dengue, chikungunya and Zika before and after differentiation. Brazil, January 2015 to December 2017.**

**S2 Table. AIC and Lag values for the restricts and unrestricted models for the series confirmed and discarded cases of dengue, chikungunya and Zika. Brazil, January 2015 to December 2017.**

**S3 Table. Results of pairwise Granger tests for restricted models.** Exploratory search of associations between series of confirmed and discarded cases of dengue, chikungunya and Zika. Brazil, January 2015 to December 2017.

**S1 Fig. Autocorrelation function (ACF) plot of the residuals with** 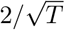 **bounds.** The plots along the diagonal are the individual ACFs for each model’s residuals, the remaining subplots show the cross-correlations between pairwise residuals. Here *T* is the sample size. The plots suggests randomness of the residuals, indicating a good fitting of the VAR model.

**S2 Fig. Probability plot and histograms for confirmed and discarded Zika model’s residuals.** The probability plot shows the unscaled quantiles of residuals versus the probabilities of a normal distribution.

**S3 Fig. Probability plot and histograms for confirmed and discarded chikungunya model’s residuals.** The probability plot shows the unscaled quantiles of residuals versus the probabilities of a normal distribution.

**S4 Fig. Probability plot and histograms for confirmed and discarded dengue model’s residuals.** The probability plot shows the unscaled quantiles of residuals versus the probabilities of a normal distribution.

**S1 Appendix. Summary of regression results.** VAR model for the 6-dimensional multivariate time series analysis of confirmed and discarded cases of dengue, chikungunya and Zika.

**S2 Appendix.** STROBE Check-list

## Acknowledgments

We would like to thank all the specialists for the discussions and support regarding the modeling approach.

