## Supplementary material for "Interdependence between confirmed and discarded cases of dengue, chikungunya and Zika viruses in Brazil: A multivariate time-series analysis": Results of Dickey-Fuller Test applied to the series of confirmed and discarded cases of dengue, chikungunya and Zika before and after differentiation.

Table 1: Results of Dickey-Fuller Test applied to the series of confirmed and discarded cases of dengue, chikungunya and Zika before and after differentiation. Brazil, January 2015 to December 2017.

| | $Z_{1t}$ | $Z_{2t}$ | $C_{1t}$ | $C_{2t}$ | $D_{2t}$ | $D_{2t}$ | $DZ_{1t}$ | $DZ_{2t}$ | $DC_{1t}$ | $DC_{2t}$ | $DD_{2t}$ | $DD_{2t}$ |
|---|---|---|---|---|---|---|---|---|---|---|---|---|
| Test Statistic | -2.25 | -2.33 | -2.36 | -1.90 | -4.29 | -2.60 | -3.68 | -4.50 | -4.92 | -6.89 | -5.34 | -5.58 |
| p-value | 0.19 | 0.16 | 0.15 | 0.33 | $<0.001$ | 0.09 | 0.004 | $<0.001$ | $<0.001$ | $<0.001$ | $<0.001$ | $<0.001$ |
| #Lags Used | 12 | 8 | 6 | 2 | 6 | 10 | 11 | 7 | 5 | 1 | 10 | 9 |
| Number of Observations Used | 143 | 147 | 149 | 153 | 149 | 145 | 143 | 147 | 149 | 153 | 144 | 145 |
