## Supplementary material for "Interdependence between confirmed and discarded cases of dengue, chikungunya and Zika viruses in Brazil: A multivariate time-series analysis": AIC and Lag values for the restricts and unrestricted models for the series confirmed and discarded cases of dengue, chikungunya and Zika. Brazil, Jan

Table 1: AIC and Lag values for the restricts and unrestricted models for the series confirmed and discarded cases of dengue, chikungunya and Zika. Brazil, January 2015 to December 2017.

| | Unrestricted model | Restricted model without Zika series | Restricted model without chikungunya series | Restricted model without Dengue series |
|---|---|---|---|---|
| AIC values | 69.44 | 52.06 | 48.50 | 41.67 |
| #Lags | 13 | 12 | 13 | 12 |
