## Supplementary material for "Interdependence between confirmed and discarded cases of dengue, chikungunya and Zika viruses in Brazil: A multivariate time-series analysis": Results of pairwise Granger tests for restricted models.

**Table 1. Results of pairwise Granger tests for restricted models.** Exploratory search of associations between series of confirmed and discarded cases of dengue, chikungunya and Zika. Brazil, January 2015 to December 2017.

| Restricted model without Zika series | | | |
|---|---|---|---|
| Null hypothesis | Test statistic | p-value | Result |
| Discarded cases of chikungunya do not affect confirmed cases of dengue | 2.284 | 0.008 | Reject |
| Confirmed cases of dengue do not affect confirmed cases of chikungunya | 0.7111 | 0.74 | Do not reject |
| Confirmed cases of chikungunya do not affect confirmed cases of dengue | 1.300 | 0.216 | Do not reject |
| Confirmed cases of dengue do not affect discarded cases of chikungunya | 0.9936 | 0.454 | Do not reject |
| Discarded cases of dengue do not affect confirmed cases of chikungunya | 0.7065 | 0.745 | Do not reject |
| Confirmed cases of chikungunya do not affect discarded cases of dengue | 1.417 | 0.155 | Do not reject |
| Restricted model without dengue series | | | |
| Confirmed cases of Zika do not affect discarded cases of chikungunya | 2.206 | 0.011 | Reject |
| Discarded cases of Zika do not affect confirmed cases of chikungunya | 2.447 | 0.004 | Reject |
| Confirmed cases of Zika do not affect confirmed chikungunya | 1.761 | 0.053 | Do not reject |
| Confirmed cases of chikungunya do not affect confirmed cases of Zika | 0.7291 | 0.723 | Do not reject |
| Discarded cases of chikungunya do not affect confirmed cases of Zika | 1.016 | 0.433 | Do not reject |
| Discarded cases of Zika do not affect confirmed cases of chikungunya | 0.8772 | 0.571 | Do not reject |
| Restricted model without chikungunya series | | | |
| Confirmed cases of Zika do not affect discarded cases of dengue | 4.377 | $< 0.001$ | Reject |
| Discarded cases of dengue do not affect confirmed cases of Zika | 5.227 | $< 0.001$ | Reject |
| Confirmed cases of dengue do not affect discarded cases of Zika | 5.479 | $< 0.001$ | Reject |
| Confirmed cases of Zika do not affect confirmed cases of dengue | 6.442 | $< 0.001$ | Reject |
| Confirmed cases of dengue do not affect confirmed cases of Zika | 5.227 | $< 0.001$ | Reject |
| Discarded cases of Zika do not affect confirmed cases of dengue | 4.690 | $< 0.001$ | Reject |
