## Supplementary material for "Interdependence between confirmed and discarded cases of dengue, chikungunya and Zika viruses in Brazil: A multivariate time-series analysis": Summary of regression results

| | |
|---|---|
| Model: | VAR |
| Method: | OLS |
| Date: | Thu, 26, Dec, 2019 |
| Time: | 12:48:20 |

| | | | |
|---|---|---|---|
| No. of Equations: | 6.00000 | BIC: | 79.3113 |
| Nobs: | 142.000 | HQIC: | 73.4541 |
| Log likelihood: | -5665.51 | FPE: | 3.39136e+30 |
| AIC: | 69.4447 | Det(Omega_mle): | 2.38644e+29 |

Results for equation cases_zika

| | coefficient | std. error | t-stat | prob |
|---|---|---|---|---|
| const | 11.094524 | 28.124098 | 0.394 | 0.693 |
| L1.cases_zika | 0.897923 | 0.168115 | 5.341 | 0.000 |
| L1.cases_des_zika | 1.533060 | 0.730488 | 2.099 | 0.036 |
| L1.cases_chik | 0.048381 | 0.149485 | 0.324 | 0.746 |
| L1.cases_des_chik | -0.298380 | 0.406153 | -0.735 | 0.463 |
| L1.cases_dengue | -0.046167 | 0.021204 | -2.177 | 0.029 |
| L1.cases_dengue_des | -0.158328 | 0.049182 | -3.219 | 0.001 |
| L2.cases_zika | -0.690075 | 0.203275 | -3.395 | 0.001 |
| L2.cases_des_zika | 0.943165 | 0.792161 | 1.191 | 0.234 |
| L2.cases_chik | 0.020578 | 0.137753 | 0.149 | 0.881 |
| L2.cases_des_chik | 0.017230 | 0.411329 | 0.042 | 0.967 |
| L2.cases_dengue | 0.136659 | 0.025724 | 5.312 | 0.000 |
| L2.cases_dengue_des | -0.084641 | 0.057561 | -1.470 | 0.141 |
| L3.cases_zika | -0.211701 | 0.228697 | -0.926 | 0.355 |
| L3.cases_des_zika | -0.213419 | 0.762746 | -0.280 | 0.780 |
| L3.cases_chik | -0.153490 | 0.160796 | -0.955 | 0.340 |
| L3.cases_des_chik | 0.854566 | 0.443824 | 1.925 | 0.054 |
| L3.cases_dengue | -0.018555 | 0.031249 | -0.594 | 0.553 |
| L3.cases_dengue_des | 0.085785 | 0.062131 | 1.381 | 0.167 |
| L4.cases_zika | 0.036012 | 0.234096 | 0.154 | 0.878 |
| L4.cases_des_zika | 0.413401 | 0.868047 | 0.476 | 0.634 |
| L4.cases_chik | -0.172107 | 0.152852 | -1.126 | 0.260 |
| L4.cases_des_chik | -0.413619 | 0.462932 | -0.893 | 0.372 |
| L4.cases_dengue | -0.015609 | 0.028930 | -0.540 | 0.589 |
| L4.cases_dengue_des | 0.071332 | 0.061799 | 1.154 | 0.248 |
| L5.cases_zika | 0.201610 | 0.209063 | 0.964 | 0.335 |
| L5.cases_des_zika | 0.165454 | 0.822154 | 0.201 | 0.841 |
| L5.cases_chik | 0.072781 | 0.154958 | 0.470 | 0.639 |
| L5.cases_des_chik | -0.189762 | 0.459919 | -0.413 | 0.680 |
| L5.cases_dengue | 0.015424 | 0.029177 | 0.529 | 0.597 |
| L5.cases_dengue_des | 0.002881 | 0.052582 | 0.055 | 0.956 |
| L6.cases_zika | -0.123935 | 0.227611 | -0.545 | 0.586 |
| L6.cases_des_zika | -1.456457 | 0.888872 | -1.639 | 0.101 |
| L6.cases_chik | 0.091341 | 0.161122 | 0.567 | 0.571 |
| L6.cases_des_chik | -0.271269 | 0.449494 | -0.603 | 0.546 |
| L6.cases_dengue | 0.050604 | 0.024518 | 2.064 | 0.039 |
| L6.cases_dengue_des | 0.168810 | 0.046286 | 3.647 | 0.000 |

| | | | | |
|---|---|---|---|---|
| L7.cases_zika | -0.417545 | 0.213913 | -1.952 | 0.051 |
| L7.cases_des_zika | 1.985676 | 0.834852 | 2.378 | 0.017 |
| L7.cases_chik | -0.091553 | 0.169917 | -0.539 | 0.590 |
| L7.cases_des_chik | -0.339353 | 0.449194 | -0.755 | 0.450 |
| L7.cases_dengue | -0.168528 | 0.026585 | -6.339 | 0.000 |
| L7.cases_dengue_des | 0.053466 | 0.044131 | 1.212 | 0.226 |
| L8.cases_zika | 0.304054 | 0.214296 | 1.419 | 0.156 |
| L8.cases_des_zika | 0.149818 | 0.755352 | 0.198 | 0.843 |
| L8.cases_chik | 0.107014 | 0.158030 | 0.677 | 0.498 |
| L8.cases_des_chik | -0.164668 | 0.443811 | -0.371 | 0.711 |
| L8.cases_dengue | 0.051833 | 0.029626 | 1.750 | 0.080 |
| L8.cases_dengue_des | -0.033110 | 0.047213 | -0.701 | 0.483 |
| L9.cases_zika | 0.054633 | 0.209230 | 0.261 | 0.794 |
| L9.cases_des_zika | -0.967618 | 0.764804 | -1.265 | 0.206 |
| L9.cases_chik | 0.226059 | 0.156001 | 1.449 | 0.147 |
| L9.cases_des_chik | -0.424262 | 0.452251 | -0.938 | 0.348 |
| L9.cases_dengue | -0.025822 | 0.030130 | -0.857 | 0.391 |
| L9.cases_dengue_des | 0.022919 | 0.049982 | 0.459 | 0.647 |
| L10.cases_zika | -0.080753 | 0.202077 | -0.400 | 0.689 |
| L10.cases_des_zika | -1.430709 | 0.727667 | -1.966 | 0.049 |
| L10.cases_chik | -0.225482 | 0.163548 | -1.379 | 0.168 |
| L10.cases_des_chik | 0.550342 | 0.442219 | 1.245 | 0.213 |
| L10.cases_dengue | -0.036137 | 0.028367 | -1.274 | 0.203 |
| L10.cases_dengue_des | 0.098266 | 0.047316 | 2.077 | 0.038 |
| L11.cases_zika | 0.628872 | 0.206281 | 3.049 | 0.002 |
| L11.cases_des_zika | -1.458785 | 0.784026 | -1.861 | 0.063 |
| L11.cases_chik | 0.078187 | 0.163313 | 0.479 | 0.632 |
| L11.cases_des_chik | -0.397749 | 0.447309 | -0.889 | 0.374 |
| L11.cases_dengue | -0.053401 | 0.027257 | -1.959 | 0.050 |
| L11.cases_dengue_des | 0.030319 | 0.045203 | 0.671 | 0.502 |
| L12.cases_zika | -0.401798 | 0.208689 | -1.925 | 0.054 |
| L12.cases_des_zika | 1.027986 | 0.840143 | 1.224 | 0.221 |
| L12.cases_chik | -0.031798 | 0.151355 | -0.210 | 0.834 |
| L12.cases_des_chik | 0.072964 | 0.412643 | 0.177 | 0.860 |
| L12.cases_dengue | 0.035607 | 0.026831 | 1.327 | 0.184 |
| L12.cases_dengue_des | 0.044942 | 0.043858 | 1.025 | 0.306 |
| L13.cases_zika | -0.197608 | 0.211541 | -0.934 | 0.350 |
| L13.cases_des_zika | -0.200851 | 0.862065 | -0.233 | 0.816 |
| L13.cases_chik | 0.064924 | 0.158917 | 0.409 | 0.683 |
| L13.cases_des_chik | -0.071529 | 0.409352 | -0.175 | 0.861 |
| L13.cases_dengue | -0.043492 | 0.026734 | -1.627 | 0.104 |
| L13.cases_dengue_des | 0.110564 | 0.034900 | 3.168 | 0.002 |

Results for equation cases_des_zika

| | coefficient | std. error | t-stat | prob |
|---|---|---|---|---|
| const | 3.500321 | 6.695504 | 0.523 | 0.601 |
| L1.cases_zika | 0.153770 | 0.040023 | 3.842 | 0.000 |
| L1.cases_des_zika | 0.545798 | 0.173907 | 3.138 | 0.002 |
| L1.cases_chik | -0.002482 | 0.035588 | -0.070 | 0.944 |
| L1.cases_des_chik | -0.016342 | 0.096693 | -0.169 | 0.866 |

| | | | | |
|---|---|---|---|---|
| L1.cases_dengue | -0.000862 | 0.005048 | -0.171 | 0.864 |
| L1.cases_dengue_des | -0.058296 | 0.011709 | -4.979 | 0.000 |
| L2.cases_zika | -0.098163 | 0.048394 | -2.028 | 0.043 |
| L2.cases_des_zika | 0.195727 | 0.188590 | 1.038 | 0.299 |
| L2.cases_chik | 0.055188 | 0.032795 | 1.683 | 0.092 |
| L2.cases_des_chik | -0.063797 | 0.097925 | -0.651 | 0.515 |
| L2.cases_dengue | 0.027516 | 0.006124 | 4.493 | 0.000 |
| L2.cases_dengue_des | -0.027617 | 0.013703 | -2.015 | 0.044 |
| L3.cases_zika | -0.104338 | 0.054446 | -1.916 | 0.055 |
| L3.cases_des_zika | 0.323224 | 0.181587 | 1.780 | 0.075 |
| L3.cases_chik | -0.045744 | 0.038281 | -1.195 | 0.232 |
| L3.cases_des_chik | 0.209283 | 0.105661 | 1.981 | 0.048 |
| L3.cases_dengue | 0.002302 | 0.007439 | 0.309 | 0.757 |
| L3.cases_dengue_des | 0.002193 | 0.014791 | 0.148 | 0.882 |
| L4.cases_zika | 0.041437 | 0.055731 | 0.744 | 0.457 |
| L4.cases_des_zika | -0.143058 | 0.206656 | -0.692 | 0.489 |
| L4.cases_chik | -0.067515 | 0.036390 | -1.855 | 0.064 |
| L4.cases_des_chik | 0.045655 | 0.110210 | 0.414 | 0.679 |
| L4.cases_dengue | -0.000892 | 0.006887 | -0.129 | 0.897 |
| L4.cases_dengue_des | 0.025749 | 0.014713 | 1.750 | 0.080 |
| L5.cases_zika | 0.084788 | 0.049772 | 1.704 | 0.088 |
| L5.cases_des_zika | -0.365668 | 0.195730 | -1.868 | 0.062 |
| L5.cases_chik | -0.002845 | 0.036891 | -0.077 | 0.939 |
| L5.cases_des_chik | -0.031000 | 0.109493 | -0.283 | 0.777 |
| L5.cases_dengue | 0.002020 | 0.006946 | 0.291 | 0.771 |
| L5.cases_dengue_des | 0.016140 | 0.012518 | 1.289 | 0.197 |
| L6.cases_zika | 0.035522 | 0.054187 | 0.656 | 0.512 |
| L6.cases_des_zika | -0.221049 | 0.211614 | -1.045 | 0.296 |
| L6.cases_chik | -0.007814 | 0.038358 | -0.204 | 0.839 |
| L6.cases_des_chik | -0.052905 | 0.107011 | -0.494 | 0.621 |
| L6.cases_dengue | 0.005391 | 0.005837 | 0.924 | 0.356 |
| L6.cases_dengue_des | 0.034974 | 0.011019 | 3.174 | 0.002 |
| L7.cases_zika | -0.074749 | 0.050926 | -1.468 | 0.142 |
| L7.cases_des_zika | 0.212385 | 0.198753 | 1.069 | 0.285 |
| L7.cases_chik | 0.022033 | 0.040452 | 0.545 | 0.586 |
| L7.cases_des_chik | -0.147372 | 0.106940 | -1.378 | 0.168 |
| L7.cases_dengue | -0.026251 | 0.006329 | -4.148 | 0.000 |
| L7.cases_dengue_des | 0.024216 | 0.010506 | 2.305 | 0.021 |
| L8.cases_zika | 0.048191 | 0.051017 | 0.945 | 0.345 |
| L8.cases_des_zika | 0.060713 | 0.179827 | 0.338 | 0.736 |
| L8.cases_chik | 0.022819 | 0.037622 | 0.607 | 0.544 |
| L8.cases_des_chik | 0.050648 | 0.105658 | 0.479 | 0.632 |
| L8.cases_dengue | 0.001218 | 0.007053 | 0.173 | 0.863 |
| L8.cases_dengue_des | -0.003618 | 0.011240 | -0.322 | 0.748 |
| L9.cases_zika | 0.011073 | 0.049811 | 0.222 | 0.824 |
| L9.cases_des_zika | -0.209837 | 0.182077 | -1.152 | 0.249 |
| L9.cases_chik | 0.027751 | 0.037139 | 0.747 | 0.455 |
| L9.cases_des_chik | -0.024009 | 0.107668 | -0.223 | 0.824 |
| L9.cases_dengue | -0.005489 | 0.007173 | -0.765 | 0.444 |
| L9.cases_dengue_des | 0.003254 | 0.011899 | 0.273 | 0.785 |
| L10.cases_zika | 0.064279 | 0.048109 | 1.336 | 0.182 |
| L10.cases_des_zika | -0.590941 | 0.173236 | -3.411 | 0.001 |
| L10.cases_chik | -0.016240 | 0.038936 | -0.417 | 0.677 |

| | | | | |
|---|---|---|---|---|
| L10.cases_des_chik | 0.115992 | 0.105279 | 1.102 | 0.271 |
| L10.cases_dengue | -0.009162 | 0.006753 | -1.357 | 0.175 |
| L10.cases_dengue_des | 0.017654 | 0.011265 | 1.567 | 0.117 |
| L11.cases_zika | 0.186632 | 0.049109 | 3.800 | 0.000 |
| L11.cases_des_zika | -0.662448 | 0.186653 | -3.549 | 0.000 |
| L11.cases_chik | 0.009343 | 0.038880 | 0.240 | 0.810 |
| L11.cases_des_chik | -0.042072 | 0.106491 | -0.395 | 0.693 |
| L11.cases_dengue | -0.010161 | 0.006489 | -1.566 | 0.117 |
| L11.cases_dengue_des | 0.018022 | 0.010761 | 1.675 | 0.094 |
| L12.cases_zika | -0.116973 | 0.049683 | -2.354 | 0.019 |
| L12.cases_des_zika | 0.238299 | 0.200013 | 1.191 | 0.233 |
| L12.cases_chik | -0.032113 | 0.036033 | -0.891 | 0.373 |
| L12.cases_des_chik | 0.009633 | 0.098238 | 0.098 | 0.922 |
| L12.cases_dengue | -0.004770 | 0.006388 | -0.747 | 0.455 |
| L12.cases_dengue_des | 0.022606 | 0.010441 | 2.165 | 0.030 |
| L13.cases_zika | -0.002168 | 0.050362 | -0.043 | 0.966 |
| L13.cases_des_zika | -0.070744 | 0.205232 | -0.345 | 0.730 |
| L13.cases_chik | -0.009417 | 0.037833 | -0.249 | 0.803 |
| L13.cases_des_chik | 0.106187 | 0.097454 | 1.090 | 0.276 |
| L13.cases_dengue | -0.005714 | 0.006365 | -0.898 | 0.369 |
| L13.cases_dengue_des | 0.013796 | 0.008309 | 1.660 | 0.097 |

Results for equation cases_chik

| | coefficient | std. error | t-stat | prob |
|---|---|---|---|---|
| const | -13.915982 | 27.337543 | -0.509 | 0.611 |
| L1.cases_zika | -0.184642 | 0.163413 | -1.130 | 0.259 |
| L1.cases_des_zika | 0.680923 | 0.710058 | 0.959 | 0.338 |
| L1.cases_chik | -0.070040 | 0.145304 | -0.482 | 0.630 |
| L1.cases_des_chik | -0.651946 | 0.394794 | -1.651 | 0.099 |
| L1.cases_dengue | 0.000021 | 0.020611 | 0.001 | 0.999 |
| L1.cases_dengue_des | 0.051685 | 0.047806 | 1.081 | 0.280 |
| L2.cases_zika | -0.074540 | 0.197590 | -0.377 | 0.706 |
| L2.cases_des_zika | 1.260483 | 0.770006 | 1.637 | 0.102 |
| L2.cases_chik | 0.712833 | 0.133901 | 5.324 | 0.000 |
| L2.cases_des_chik | -1.693991 | 0.399825 | -4.237 | 0.000 |
| L2.cases_dengue | -0.050748 | 0.025005 | -2.030 | 0.042 |
| L2.cases_dengue_des | 0.074683 | 0.055951 | 1.335 | 0.182 |
| L3.cases_zika | -0.457288 | 0.222300 | -2.057 | 0.040 |
| L3.cases_des_zika | 0.872903 | 0.741414 | 1.177 | 0.239 |
| L3.cases_chik | 0.374834 | 0.156299 | 2.398 | 0.016 |
| L3.cases_des_chik | -1.122729 | 0.431412 | -2.602 | 0.009 |
| L3.cases_dengue | -0.009599 | 0.030375 | -0.316 | 0.752 |
| L3.cases_dengue_des | 0.117891 | 0.060393 | 1.952 | 0.051 |
| L4.cases_zika | 0.180660 | 0.227549 | 0.794 | 0.427 |
| L4.cases_des_zika | -1.596248 | 0.843770 | -1.892 | 0.059 |
| L4.cases_chik | -0.134500 | 0.148577 | -0.905 | 0.365 |
| L4.cases_des_chik | 0.969467 | 0.449985 | 2.154 | 0.031 |
| L4.cases_dengue | -0.028074 | 0.028121 | -0.998 | 0.318 |
| L4.cases_dengue_des | 0.098683 | 0.060071 | 1.643 | 0.100 |
| L5.cases_zika | -0.432976 | 0.203216 | -2.131 | 0.033 |

| | | | | |
|---|---|---|---|---|
| L5.cases_des_zika | 0.181582 | 0.799161 | 0.227 | 0.820 |
| L5.cases_chik | 0.090081 | 0.150624 | 0.598 | 0.550 |
| L5.cases_des_chik | -0.761246 | 0.447056 | -1.703 | 0.089 |
| L5.cases_dengue | -0.015916 | 0.028361 | -0.561 | 0.575 |
| L5.cases_dengue_des | 0.121323 | 0.051111 | 2.374 | 0.018 |
| L6.cases_zika | 0.207648 | 0.221246 | 0.939 | 0.348 |
| L6.cases_des_zika | 0.245591 | 0.864013 | 0.284 | 0.776 |
| L6.cases_chik | -0.436098 | 0.156616 | -2.785 | 0.005 |
| L6.cases_des_chik | 0.605027 | 0.436923 | 1.385 | 0.166 |
| L6.cases_dengue | -0.074590 | 0.023832 | -3.130 | 0.002 |
| L6.cases_dengue_des | 0.103928 | 0.044992 | 2.310 | 0.021 |
| L7.cases_zika | -0.364973 | 0.207931 | -1.755 | 0.079 |
| L7.cases_des_zika | 0.416176 | 0.811503 | 0.513 | 0.608 |
| L7.cases_chik | 0.247243 | 0.165165 | 1.497 | 0.134 |
| L7.cases_des_chik | -0.358135 | 0.436632 | -0.820 | 0.412 |
| L7.cases_dengue | -0.014547 | 0.025841 | -0.563 | 0.573 |
| L7.cases_dengue_des | 0.096209 | 0.042897 | 2.243 | 0.025 |
| L8.cases_zika | -0.103750 | 0.208303 | -0.498 | 0.618 |
| L8.cases_des_zika | -0.667996 | 0.734227 | -0.910 | 0.363 |
| L8.cases_chik | -0.269907 | 0.153610 | -1.757 | 0.079 |
| L8.cases_des_chik | 1.534154 | 0.431399 | 3.556 | 0.000 |
| L8.cases_dengue | -0.068830 | 0.028797 | -2.390 | 0.017 |
| L8.cases_dengue_des | 0.164204 | 0.045893 | 3.578 | 0.000 |
| L9.cases_zika | -0.001881 | 0.203378 | -0.009 | 0.993 |
| L9.cases_des_zika | -0.436446 | 0.743415 | -0.587 | 0.557 |
| L9.cases_chik | -0.214708 | 0.151638 | -1.416 | 0.157 |
| L9.cases_des_chik | 0.915687 | 0.439603 | 2.083 | 0.037 |
| L9.cases_dengue | -0.066114 | 0.029288 | -2.257 | 0.024 |
| L9.cases_dengue_des | 0.115287 | 0.048584 | 2.373 | 0.018 |
| L10.cases_zika | 0.150883 | 0.196426 | 0.768 | 0.442 |
| L10.cases_des_zika | -0.380277 | 0.707316 | -0.538 | 0.591 |
| L10.cases_chik | 0.098955 | 0.158974 | 0.622 | 0.534 |
| L10.cases_des_chik | 0.897270 | 0.429851 | 2.087 | 0.037 |
| L10.cases_dengue | -0.050719 | 0.027574 | -1.839 | 0.066 |
| L10.cases_dengue_des | 0.097272 | 0.045993 | 2.115 | 0.034 |
| L11.cases_zika | -0.112929 | 0.200512 | -0.563 | 0.573 |
| L11.cases_des_zika | -0.069715 | 0.762098 | -0.091 | 0.927 |
| L11.cases_chik | 0.098362 | 0.158746 | 0.620 | 0.536 |
| L11.cases_des_chik | -0.245954 | 0.434799 | -0.566 | 0.572 |
| L11.cases_dengue | -0.038058 | 0.026494 | -1.436 | 0.151 |
| L11.cases_dengue_des | 0.129383 | 0.043939 | 2.945 | 0.003 |
| L12.cases_zika | -0.306173 | 0.202853 | -1.509 | 0.131 |
| L12.cases_des_zika | 0.752396 | 0.816646 | 0.921 | 0.357 |
| L12.cases_chik | -0.419601 | 0.147122 | -2.852 | 0.004 |
| L12.cases_des_chik | 0.833122 | 0.401102 | 2.077 | 0.038 |
| L12.cases_dengue | -0.075604 | 0.026081 | -2.899 | 0.004 |
| L12.cases_dengue_des | 0.092518 | 0.042632 | 2.170 | 0.030 |
| L13.cases_zika | 0.114706 | 0.205625 | 0.558 | 0.577 |
| L13.cases_des_zika | -0.213096 | 0.837955 | -0.254 | 0.799 |
| L13.cases_chik | -0.236499 | 0.154472 | -1.531 | 0.126 |
| L13.cases_des_chik | 0.575234 | 0.397904 | 1.446 | 0.148 |
| L13.cases_dengue | -0.010782 | 0.025987 | -0.415 | 0.678 |
| L13.cases_dengue_des | 0.023736 | 0.033924 | 0.700 | 0.484 |

==============================================================================

Results for equation cases_des_chik

| | coefficient | std. error | t-stat | prob |
|---|---|---|---|---|
| const | 1.283177 | 11.173407 | 0.115 | 0.909 |
| L1.cases_zika | -0.035587 | 0.066790 | -0.533 | 0.594 |
| L1.cases_des_zika | 0.381725 | 0.290215 | 1.315 | 0.188 |
| L1.cases_chik | -0.063587 | 0.059389 | -1.071 | 0.284 |
| L1.cases_des_chik | -0.042706 | 0.161360 | -0.265 | 0.791 |
| L1.cases_dengue | 0.005938 | 0.008424 | 0.705 | 0.481 |
| L1.cases_dengue_des | -0.015892 | 0.019539 | -0.813 | 0.416 |
| L2.cases_zika | -0.094376 | 0.080759 | -1.169 | 0.243 |
| L2.cases_des_zika | 0.244900 | 0.314717 | 0.778 | 0.436 |
| L2.cases_chik | 0.105473 | 0.054728 | 1.927 | 0.054 |
| L2.cases_des_chik | -0.266636 | 0.163417 | -1.632 | 0.103 |
| L2.cases_dengue | -0.003935 | 0.010220 | -0.385 | 0.700 |
| L2.cases_dengue_des | -0.001426 | 0.022868 | -0.062 | 0.950 |
| L3.cases_zika | -0.125927 | 0.090859 | -1.386 | 0.166 |
| L3.cases_des_zika | 0.880798 | 0.303031 | 2.907 | 0.004 |
| L3.cases_chik | 0.002108 | 0.063882 | 0.033 | 0.974 |
| L3.cases_des_chik | -0.468757 | 0.176327 | -2.658 | 0.008 |
| L3.cases_dengue | 0.000310 | 0.012415 | 0.025 | 0.980 |
| L3.cases_dengue_des | 0.014819 | 0.024684 | 0.600 | 0.548 |
| L4.cases_zika | 0.035549 | 0.093004 | 0.382 | 0.702 |
| L4.cases_des_zika | -0.268716 | 0.344866 | -0.779 | 0.436 |
| L4.cases_chik | -0.016094 | 0.060727 | -0.265 | 0.791 |
| L4.cases_des_chik | -0.063439 | 0.183918 | -0.345 | 0.730 |
| L4.cases_dengue | -0.003803 | 0.011493 | -0.331 | 0.741 |
| L4.cases_dengue_des | 0.039624 | 0.024552 | 1.614 | 0.107 |
| L5.cases_zika | -0.136398 | 0.083059 | -1.642 | 0.101 |
| L5.cases_des_zika | -0.495480 | 0.326633 | -1.517 | 0.129 |
| L5.cases_chik | 0.193498 | 0.061563 | 3.143 | 0.002 |
| L5.cases_des_chik | -0.398090 | 0.182721 | -2.179 | 0.029 |
| L5.cases_dengue | 0.000434 | 0.011592 | 0.037 | 0.970 |
| L5.cases_dengue_des | 0.071603 | 0.020890 | 3.428 | 0.001 |
| L6.cases_zika | 0.090106 | 0.090428 | 0.996 | 0.319 |
| L6.cases_des_zika | 0.006465 | 0.353140 | 0.018 | 0.985 |
| L6.cases_chik | -0.041019 | 0.064012 | -0.641 | 0.522 |
| L6.cases_des_chik | -0.000499 | 0.178579 | -0.003 | 0.998 |
| L6.cases_dengue | -0.038489 | 0.009741 | -3.951 | 0.000 |
| L6.cases_dengue_des | 0.053060 | 0.018389 | 2.885 | 0.004 |
| L7.cases_zika | -0.018830 | 0.084985 | -0.222 | 0.825 |
| L7.cases_des_zika | 0.003748 | 0.331678 | 0.011 | 0.991 |
| L7.cases_chik | 0.069117 | 0.067506 | 1.024 | 0.306 |
| L7.cases_des_chik | -0.130023 | 0.178460 | -0.729 | 0.466 |
| L7.cases_dengue | -0.007088 | 0.010562 | -0.671 | 0.502 |
| L7.cases_dengue_des | 0.012797 | 0.017533 | 0.730 | 0.465 |
| L8.cases_zika | -0.046181 | 0.085137 | -0.542 | 0.588 |
| L8.cases_des_zika | 0.101719 | 0.300094 | 0.339 | 0.735 |
| L8.cases_chik | -0.005318 | 0.062784 | -0.085 | 0.932 |
| L8.cases_des_chik | 0.366547 | 0.176322 | 2.079 | 0.038 |

| | | | | |
|---|---|---|---|---|
| L8.cases_dengue | -0.007598 | 0.011770 | -0.646 | 0.519 |
| L8.cases_dengue_des | 0.032905 | 0.018757 | 1.754 | 0.079 |
| L9.cases_zika | -0.087432 | 0.083125 | -1.052 | 0.293 |
| L9.cases_des_zika | -0.114796 | 0.303849 | -0.378 | 0.706 |
| L9.cases_chik | -0.067650 | 0.061978 | -1.092 | 0.275 |
| L9.cases_des_chik | 0.267229 | 0.179675 | 1.487 | 0.137 |
| L9.cases_dengue | -0.028796 | 0.011970 | -2.406 | 0.016 |
| L9.cases_dengue_des | 0.048682 | 0.019857 | 2.452 | 0.014 |
| L10.cases_zika | 0.009379 | 0.080283 | 0.117 | 0.907 |
| L10.cases_des_zika | -0.017948 | 0.289095 | -0.062 | 0.950 |
| L10.cases_chik | -0.079265 | 0.064976 | -1.220 | 0.222 |
| L10.cases_des_chik | 0.182226 | 0.175689 | 1.037 | 0.300 |
| L10.cases_dengue | -0.024145 | 0.011270 | -2.142 | 0.032 |
| L10.cases_dengue_des | 0.031336 | 0.018798 | 1.667 | 0.096 |
| L11.cases_zika | 0.125035 | 0.081953 | 1.526 | 0.127 |
| L11.cases_des_zika | -0.517683 | 0.311485 | -1.662 | 0.097 |
| L11.cases_chik | 0.004791 | 0.064883 | 0.074 | 0.941 |
| L11.cases_des_chik | 0.050466 | 0.177711 | 0.284 | 0.776 |
| L11.cases_dengue | -0.003391 | 0.010829 | -0.313 | 0.754 |
| L11.cases_dengue_des | 0.039186 | 0.017959 | 2.182 | 0.029 |
| L12.cases_zika | -0.086502 | 0.082910 | -1.043 | 0.297 |
| L12.cases_des_zika | 0.084077 | 0.333780 | 0.252 | 0.801 |
| L12.cases_chik | -0.039633 | 0.060132 | -0.659 | 0.510 |
| L12.cases_des_chik | 0.140431 | 0.163939 | 0.857 | 0.392 |
| L12.cases_dengue | -0.025657 | 0.010660 | -2.407 | 0.016 |
| L12.cases_dengue_des | 0.039357 | 0.017424 | 2.259 | 0.024 |
| L13.cases_zika | -0.005186 | 0.084043 | -0.062 | 0.951 |
| L13.cases_des_zika | 0.213569 | 0.342489 | 0.624 | 0.533 |
| L13.cases_chik | -0.035577 | 0.063136 | -0.564 | 0.573 |
| L13.cases_des_chik | 0.090083 | 0.162631 | 0.554 | 0.580 |
| L13.cases_dengue | -0.002852 | 0.010621 | -0.269 | 0.788 |
| L13.cases_dengue_des | 0.005171 | 0.013865 | 0.373 | 0.709 |

Results for equation cases_dengue

| | coefficient | std. error | t-stat | prob |
|---|---|---|---|---|
| const | -160.671309 | 190.319635 | -0.844 | 0.399 |
| L1.cases_zika | -0.446867 | 1.137659 | -0.393 | 0.694 |
| L1.cases_des_zika | 17.070065 | 4.943314 | 3.453 | 0.001 |
| L1.cases_chik | 0.395457 | 1.011584 | 0.391 | 0.696 |
| L1.cases_des_chik | -2.921603 | 2.748492 | -1.063 | 0.288 |
| L1.cases_dengue | 0.016670 | 0.143491 | 0.116 | 0.908 |
| L1.cases_dengue_des | -0.673699 | 0.332818 | -2.024 | 0.043 |
| L2.cases_zika | -0.585493 | 1.375590 | -0.426 | 0.670 |
| L2.cases_des_zika | -5.631557 | 5.360661 | -1.051 | 0.293 |
| L2.cases_chik | -0.412694 | 0.932197 | -0.443 | 0.658 |
| L2.cases_des_chik | 2.864799 | 2.783518 | 1.029 | 0.303 |
| L2.cases_dengue | 0.887117 | 0.174081 | 5.096 | 0.000 |
| L2.cases_dengue_des | -0.571866 | 0.389521 | -1.468 | 0.142 |
| L3.cases_zika | -6.004440 | 1.547621 | -3.880 | 0.000 |
| L3.cases_des_zika | 13.362043 | 5.161604 | 2.589 | 0.010 |

| | | | | |
|---|---|---|---|---|
| L3.cases_chik | -1.186516 | 1.088126 | -1.090 | 0.276 |
| L3.cases_des_chik | -0.487387 | 3.003420 | -0.162 | 0.871 |
| L3.cases_dengue | 0.308792 | 0.211466 | 1.460 | 0.144 |
| L3.cases_dengue_des | 0.490396 | 0.420446 | 1.166 | 0.243 |
| L4.cases_zika | 0.298268 | 1.584157 | 0.188 | 0.851 |
| L4.cases_des_zika | 3.009798 | 5.874192 | 0.512 | 0.608 |
| L4.cases_chik | -0.426530 | 1.034371 | -0.412 | 0.680 |
| L4.cases_des_chik | 0.098009 | 3.132727 | 0.031 | 0.975 |
| L4.cases_dengue | -0.170064 | 0.195771 | -0.869 | 0.385 |
| L4.cases_dengue_des | 0.585632 | 0.418203 | 1.400 | 0.161 |
| L5.cases_zika | 3.951725 | 1.414760 | 2.793 | 0.005 |
| L5.cases_des_zika | -13.656467 | 5.563630 | -2.455 | 0.014 |
| L5.cases_chik | 1.583714 | 1.048621 | 1.510 | 0.131 |
| L5.cases_des_chik | -4.127235 | 3.112334 | -1.326 | 0.185 |
| L5.cases_dengue | -0.043430 | 0.197442 | -0.220 | 0.826 |
| L5.cases_dengue_des | 0.446266 | 0.355828 | 1.254 | 0.210 |
| L6.cases_zika | 0.447195 | 1.540278 | 0.290 | 0.772 |
| L6.cases_des_zika | -5.129775 | 6.015122 | -0.853 | 0.394 |
| L6.cases_chik | -0.008736 | 1.090334 | -0.008 | 0.994 |
| L6.cases_des_chik | 1.313012 | 3.041789 | 0.432 | 0.666 |
| L6.cases_dengue | 0.058863 | 0.165918 | 0.355 | 0.723 |
| L6.cases_dengue_des | 0.444011 | 0.313226 | 1.418 | 0.156 |
| L7.cases_zika | -1.817495 | 1.447579 | -1.256 | 0.209 |
| L7.cases_des_zika | 4.443436 | 5.649558 | 0.787 | 0.432 |
| L7.cases_chik | -0.113654 | 1.149852 | -0.099 | 0.921 |
| L7.cases_des_chik | -1.348182 | 3.039760 | -0.444 | 0.657 |
| L7.cases_dengue | -0.316527 | 0.179903 | -1.759 | 0.079 |
| L7.cases_dengue_des | -0.318393 | 0.298639 | -1.066 | 0.286 |
| L8.cases_zika | -0.528749 | 1.450170 | -0.365 | 0.715 |
| L8.cases_des_zika | 12.391995 | 5.111574 | 2.424 | 0.015 |
| L8.cases_chik | 0.085895 | 1.069411 | 0.080 | 0.936 |
| L8.cases_des_chik | -0.622599 | 3.003333 | -0.207 | 0.836 |
| L8.cases_dengue | 0.268022 | 0.200480 | 1.337 | 0.181 |
| L8.cases_dengue_des | -0.660131 | 0.319498 | -2.066 | 0.039 |
| L9.cases_zika | -0.333280 | 1.415886 | -0.235 | 0.814 |
| L9.cases_des_zika | 0.307700 | 5.175534 | 0.059 | 0.953 |
| L9.cases_chik | 0.356858 | 1.055681 | 0.338 | 0.735 |
| L9.cases_des_chik | 0.542103 | 3.060448 | 0.177 | 0.859 |
| L9.cases_dengue | -0.186561 | 0.203895 | -0.915 | 0.360 |
| L9.cases_dengue_des | -0.236778 | 0.338235 | -0.700 | 0.484 |
| L10.cases_zika | -0.408625 | 1.367484 | -0.299 | 0.765 |
| L10.cases_des_zika | 0.467138 | 4.924224 | 0.095 | 0.924 |
| L10.cases_chik | -0.388443 | 1.106751 | -0.351 | 0.726 |
| L10.cases_des_chik | 0.715099 | 2.992557 | 0.239 | 0.811 |
| L10.cases_dengue | -0.244566 | 0.191965 | -1.274 | 0.203 |
| L10.cases_dengue_des | -0.004361 | 0.320193 | -0.014 | 0.989 |
| L11.cases_zika | 2.647472 | 1.395934 | 1.897 | 0.058 |
| L11.cases_des_zika | -15.606905 | 5.305609 | -2.942 | 0.003 |
| L11.cases_chik | -1.134750 | 1.105162 | -1.027 | 0.305 |
| L11.cases_des_chik | 2.730199 | 3.027003 | 0.902 | 0.367 |
| L11.cases_dengue | -0.141452 | 0.184450 | -0.767 | 0.443 |
| L11.cases_dengue_des | 0.538952 | 0.305895 | 1.762 | 0.078 |
| L12.cases_zika | -4.326466 | 1.412230 | -3.064 | 0.002 |

| | | | | |
|---|---|---|---|---|
| L12.cases_des_zika | 17.269488 | 5.685362 | 3.038 | 0.002 |
| L12.cases_chik | 0.221794 | 1.024242 | 0.217 | 0.829 |
| L12.cases_des_chik | -2.929602 | 2.792410 | -1.049 | 0.294 |
| L12.cases_dengue | -0.142483 | 0.181570 | -0.785 | 0.433 |
| L12.cases_dengue_des | 0.354326 | 0.296796 | 1.194 | 0.233 |
| L13.cases_zika | -1.819899 | 1.431527 | -1.271 | 0.204 |
| L13.cases_des_zika | 4.667516 | 5.833712 | 0.800 | 0.424 |
| L13.cases_chik | 0.408421 | 1.075411 | 0.380 | 0.704 |
| L13.cases_des_chik | -1.187314 | 2.770142 | -0.429 | 0.668 |
| L13.cases_dengue | 0.094118 | 0.180914 | 0.520 | 0.603 |
| L13.cases_dengue_des | 0.495132 | 0.236171 | 2.096 | 0.036 |

Results for equation cases_dengue_des

| | coefficient | std. error | t-stat | prob |
|---|---|---|---|---|
| const | -46.353323 | 98.349311 | -0.471 | 0.637 |
| L1.cases_zika | 0.177450 | 0.587895 | 0.302 | 0.763 |
| L1.cases_des_zika | 2.444867 | 2.554500 | 0.957 | 0.339 |
| L1.cases_chik | 0.107638 | 0.522745 | 0.206 | 0.837 |
| L1.cases_des_chik | 1.639880 | 1.420307 | 1.155 | 0.248 |
| L1.cases_dengue | 0.250785 | 0.074150 | 3.382 | 0.001 |
| L1.cases_dengue_des | -0.692674 | 0.171987 | -4.027 | 0.000 |
| L2.cases_zika | 1.725036 | 0.710848 | 2.427 | 0.015 |
| L2.cases_des_zika | -0.054749 | 2.770168 | -0.020 | 0.984 |
| L2.cases_chik | -0.568567 | 0.481721 | -1.180 | 0.238 |
| L2.cases_des_chik | 1.667821 | 1.438407 | 1.159 | 0.246 |
| L2.cases_dengue | 0.273475 | 0.089958 | 3.040 | 0.002 |
| L2.cases_dengue_des | -0.745366 | 0.201288 | -3.703 | 0.000 |
| L3.cases_zika | -3.770326 | 0.799746 | -4.714 | 0.000 |
| L3.cases_des_zika | 10.966172 | 2.667304 | 4.111 | 0.000 |
| L3.cases_chik | -0.285550 | 0.562299 | -0.508 | 0.612 |
| L3.cases_des_chik | 0.085305 | 1.552043 | 0.055 | 0.956 |
| L3.cases_dengue | 0.209278 | 0.109277 | 1.915 | 0.055 |
| L3.cases_dengue_des | 0.049734 | 0.217269 | 0.229 | 0.819 |
| L4.cases_zika | 0.992174 | 0.818627 | 1.212 | 0.226 |
| L4.cases_des_zika | -0.894029 | 3.035540 | -0.295 | 0.768 |
| L4.cases_chik | -0.026598 | 0.534520 | -0.050 | 0.960 |
| L4.cases_des_chik | 0.976526 | 1.618863 | 0.603 | 0.546 |
| L4.cases_dengue | 0.087443 | 0.101166 | 0.864 | 0.387 |
| L4.cases_dengue_des | 0.177211 | 0.216110 | 0.820 | 0.412 |
| L5.cases_zika | -0.232007 | 0.731089 | -0.317 | 0.751 |
| L5.cases_des_zika | -6.327113 | 2.875054 | -2.201 | 0.028 |
| L5.cases_chik | 0.050366 | 0.541884 | 0.093 | 0.926 |
| L5.cases_des_chik | -0.541035 | 1.608326 | -0.336 | 0.737 |
| L5.cases_dengue | 0.039690 | 0.102030 | 0.389 | 0.697 |
| L5.cases_dengue_des | 0.219223 | 0.183877 | 1.192 | 0.233 |
| L6.cases_zika | 1.340859 | 0.795952 | 1.685 | 0.092 |
| L6.cases_des_zika | -0.994711 | 3.108366 | -0.320 | 0.749 |
| L6.cases_chik | -0.939093 | 0.563440 | -1.667 | 0.096 |
| L6.cases_des_chik | 2.636285 | 1.571871 | 1.677 | 0.094 |
| L6.cases_dengue | 0.014518 | 0.085740 | 0.169 | 0.866 |

| | | | | |
|---|---|---|---|---|
| L6.cases_dengue_des | 0.136015 | 0.161862 | 0.840 | 0.401 |
| L7.cases_zika | -0.764867 | 0.748049 | -1.022 | 0.307 |
| L7.cases_des_zika | 4.145726 | 2.919458 | 1.420 | 0.156 |
| L7.cases_chik | 0.191633 | 0.594196 | 0.323 | 0.747 |
| L7.cases_des_chik | -2.489073 | 1.570822 | -1.585 | 0.113 |
| L7.cases_dengue | -0.051599 | 0.092967 | -0.555 | 0.579 |
| L7.cases_dengue_des | -0.089853 | 0.154324 | -0.582 | 0.560 |
| L8.cases_zika | -0.090293 | 0.749388 | -0.120 | 0.904 |
| L8.cases_des_zika | 5.617012 | 2.641450 | 2.126 | 0.033 |
| L8.cases_chik | -0.329741 | 0.552627 | -0.597 | 0.551 |
| L8.cases_des_chik | 1.354375 | 1.551998 | 0.873 | 0.383 |
| L8.cases_dengue | -0.035550 | 0.103600 | -0.343 | 0.731 |
| L8.cases_dengue_des | -0.166102 | 0.165103 | -1.006 | 0.314 |
| L9.cases_zika | -0.976949 | 0.731671 | -1.335 | 0.182 |
| L9.cases_des_zika | -1.514698 | 2.674502 | -0.566 | 0.571 |
| L9.cases_chik | 0.959388 | 0.545532 | 1.759 | 0.079 |
| L9.cases_des_chik | -0.888185 | 1.581513 | -0.562 | 0.574 |
| L9.cases_dengue | 0.022566 | 0.105364 | 0.214 | 0.830 |
| L9.cases_dengue_des | 0.086052 | 0.174786 | 0.492 | 0.622 |
| L10.cases_zika | -0.764555 | 0.706659 | -1.082 | 0.279 |
| L10.cases_des_zika | 0.532113 | 2.544635 | 0.209 | 0.834 |
| L10.cases_chik | -0.432713 | 0.571923 | -0.757 | 0.449 |
| L10.cases_des_chik | 1.046211 | 1.546430 | 0.677 | 0.499 |
| L10.cases_dengue | -0.122769 | 0.099200 | -1.238 | 0.216 |
| L10.cases_dengue_des | -0.032263 | 0.165463 | -0.195 | 0.845 |
| L11.cases_zika | 2.179960 | 0.721361 | 3.022 | 0.003 |
| L11.cases_des_zika | -10.306922 | 2.741719 | -3.759 | 0.000 |
| L11.cases_chik | -0.429197 | 0.571102 | -0.752 | 0.452 |
| L11.cases_des_chik | 1.078881 | 1.564230 | 0.690 | 0.490 |
| L11.cases_dengue | -0.004526 | 0.095316 | -0.047 | 0.962 |
| L11.cases_dengue_des | 0.194530 | 0.158074 | 1.231 | 0.218 |
| L12.cases_zika | -1.665553 | 0.729782 | -2.282 | 0.022 |
| L12.cases_des_zika | 8.374264 | 2.937960 | 2.850 | 0.004 |
| L12.cases_chik | -0.065857 | 0.529286 | -0.124 | 0.901 |
| L12.cases_des_chik | -0.944793 | 1.443002 | -0.655 | 0.513 |
| L12.cases_dengue | -0.099250 | 0.093828 | -1.058 | 0.290 |
| L12.cases_dengue_des | 0.086088 | 0.153372 | 0.561 | 0.575 |
| L13.cases_zika | -1.138808 | 0.739754 | -1.539 | 0.124 |
| L13.cases_des_zika | 1.008716 | 3.014621 | 0.335 | 0.738 |
| L13.cases_chik | -0.396625 | 0.555728 | -0.714 | 0.475 |
| L13.cases_des_chik | 0.241946 | 1.431495 | 0.169 | 0.866 |
| L13.cases_dengue | -0.029184 | 0.093489 | -0.312 | 0.755 |
| L13.cases_dengue_des | 0.277434 | 0.122044 | 2.273 | 0.023 |
